# Effect of Variations in the Conserved Residues E371 and S359 on the Structural Dynamics of Protein Z Dependent Protease Inhibitor (ZPI): A Molecular Dynamic Simulation Study

**DOI:** 10.1101/2020.12.02.407536

**Authors:** Chellam Gayathri Subash, Suchetana Gupta, Aravind Suresh, Sanjib Senapati, Tanusree Sengupta

## Abstract

Protein Z (PZ) dependent protease inhibitor (ZPI) is a natural anticoagulant inhibiting blood coagulation proteases fXa and fXIa. Despite being a member of the serpin superfamily, it possesses unique structural features such as activation by PZ that regulate its inhibitory function. It is well known that effecient protease inhibition by serpins is dependent on the dynamics of its reactive center loop (RCL). In order to understand the RCL dynamics of ZPI, we performed Molecular Dynamics (MD) simulation on ZPI and its E371 and S359 variants located at important conserved functional sites. Unexpectedly, the RCL of E371 variants, such as E371K, E371R, and E371Q, were shown to be very stable due to compensatory interactions at the proximal end of RCL. Interestingly, RCL flexibility was shown to be enhanced in the double mutant K318-E371 due to the repulsive effect of increased negative charge on top of the breach region. Principal component analysis (PCA) coupled with residue wise interaction network (RIN) revealed correlated motion between the RCL and the PZ binding regions in the WT. Whereas the analysis revealed a loss of regulation in correlated motion between RCL and PZ binding hotspot Tyr240 in the double mutant. Additionally, the S359F and S359I mutations also resulted in increased RCL flexibility owing to the disruption of stabilizing hydrogen bonding interaction at the N terminal end of S5A. Thus, the current study proposes that the overall stabilizing interactions of S5A is a major regulator of proper loop movement of ZPI for protease inhibition and therefore its activity. The results would be beneficial to engineer activity compromised ZPI mutants as a prophylactic agent for the treatment of hemophilia.

## Introduction

Protein Z (PZ) dependent protease inhibitor (ZPI), a natural anticoagulant present in our body, is a member of the serpin (<u>ser</u>ine <u>p</u>rotease <u>in</u>hibitor) superfamily [1]. ZPI inhibits the proteases of blood coagulation such as clotting factor Xa (fXa) in a PZ dependent and factor XIa (fXIa) in a PZ independent manner. ZPI, being an inhibitory serpin, shares a highly conserved tertiary structure with the superfamily and is composed of 9 helices, 3 beta sheets (A-C), and an extended loop known as the reactive center loop (RCL) (Figure 1A) [2]. The RCL is a highly flexible loop that renders the native fold of this class of proteins to not be their most stable state, rather to be a metastable state. The flexibility of the RCL is used to enhance the binding of serpins to their cognate protease. Upon binding to its target protease, the RCL gets cleaved and is inserted as a β-strand into its own central β–sheet (β–sheet A) along with the protease attached in its acyl form to the RCL. The loop insertion leads to a massive conformational change in which the highly dynamic RCL becomes structured and the serpin molecule gains huge structural stability. Concurrently, the covalently attached protease is translocated along with the RCL to the opposite end of the serpin and becomes inactive due to distortion of its active site. Thus, ZPI function is intricately linked to its structure, specifically the RCL dynamics. The energy barrier to full loop insertion should not be too high in order to facilitate loop insertion at a rate sufficient for protease inhibition, and, on the other hand, it should also not be too low to permit the spontaneous insertion of the loop without protease inhibition to become latent. Indeed, antithrombin, another serpin that deactivates fXa, circulates in a relatively inactive form, as the RCL is restrained due to partial insertion of its hinge region in between the β strands 3A and 5A [3]. This form spontaneously converts to a hyperstable form even at ambient temperature where the top of the β sheet opens up for full insertion of RCL. The conformational mobility of serpins also renders them prone to mutation that will lead to misfolding and homo-polymerization [4].

**Figure 1.**
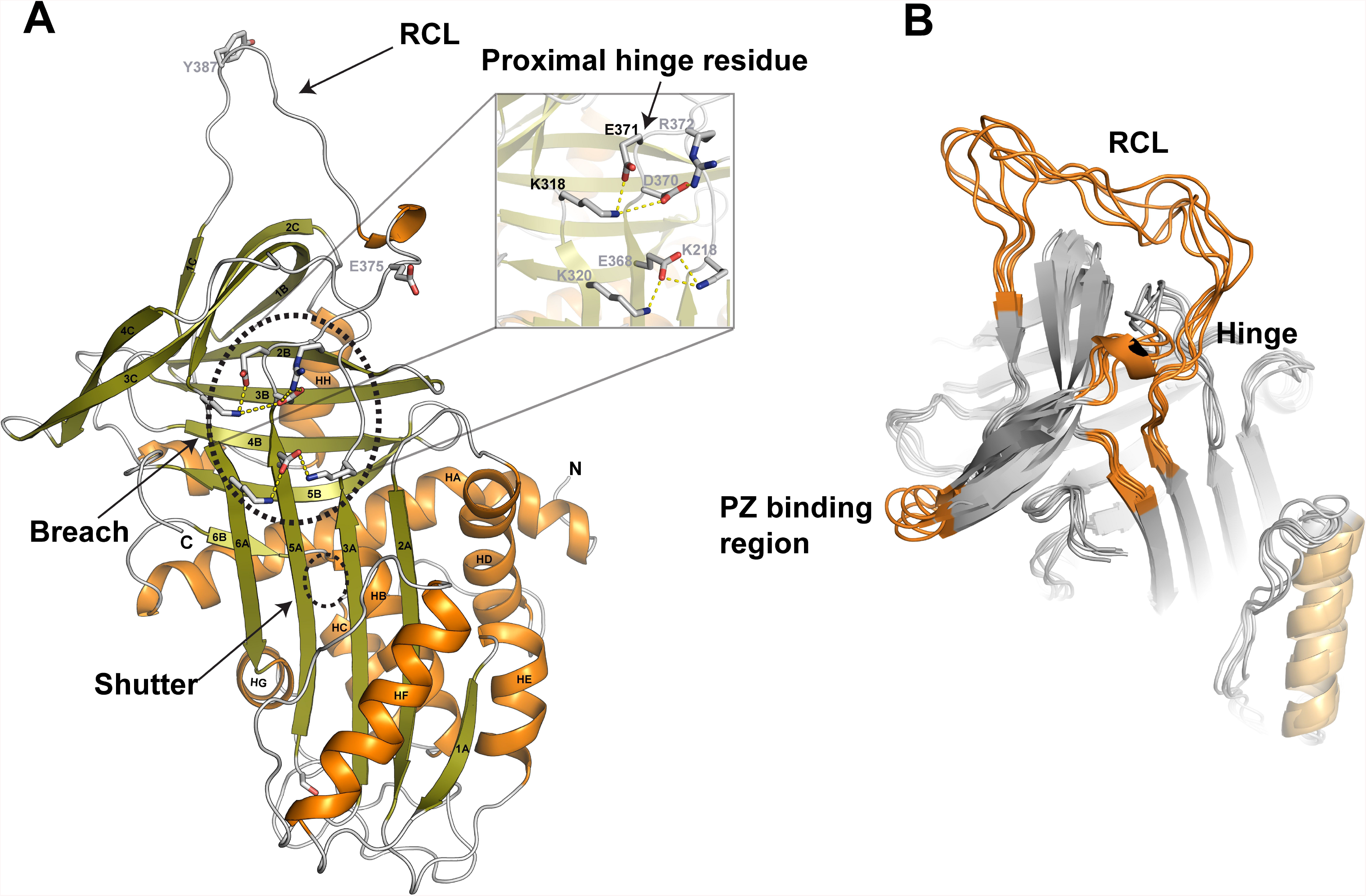
A) Structure of human ZPI (PDB ID: 3F1S) indicating the nomenclature of the helices and strands. The positions of the breach, shutter, proximal-hinge, and RCL are indicated. B) Superposition of 5 structures of the WT sampled every 10 ns from a 100-ns simulation of the WT showing regions that undergo maximum fluctuations during the simulation.

Lack of balance between the metastable active and latent or polymerized inactive conformations of serpin results in impaired protease inhibition and can lead to several physiological complications. However, in case of ZPI, such dysfunction can be beneficial as it can restore hemostasis in hemophilia patients [5,6]. In hemophilia patients, suppression of activity of the natural anticoagulants to promote clotting is a promising new approach of treatment [7–10], and the most potent candidate for this therapy is, ZPI and/ZPI-PZ complex. Low level of ZPI/ZPI-PZ is not associated with thrombotic complications as it limits coagulation only modestly, whereas, even a mild deficiency of other endogenous anticoagulant such as tissue factor pathway inhibitor (TFPI), antithrombin (AT), protein C (PC) and protein S (PS) would pose a high risk of thrombosis in hemophilia patients [6]. However, in order to engineer activity-compromised ZPI as an anti-hemophilic agent, it is absolutely pertinent to study RCL dynamics of ZPI and identify the residues controlling it. Several highly conserved residues in the hinge and shutter region of other serpins have been reported to regulate the flexibility of the RCL and consequently alter their activity [3,11]. Therefore, the effect of mutation of those conserved residues (Figure 1A, inset) on the conformational dynamics and overall stability of ZPI would be interesting to study. In turn, this would enable us to predict the residues critical for the inhibitory activity of ZPI-PZ and/ or ZPI.

Unlike other serpins, ZPI mediated protease inhibition undergoes a 1000-fold rate enhancement in the presence of its cofactor PZ, the procoagulant lipid and Ca^2+^. PZ binds to the lipid membrane through its Gla(gamma carboxy <u>gl</u>utamic <u>a</u>cid rich) domain and positions ZPI, complexed with PZ, close to its target protease (fXa) bound to the same membrane [12]. Thus unlike antithrombin, ZPI specifically inhibits fXa only if it is bound to the membrane or if fXa is bound to its cofactor fVa in prothrombinase complex [13]. Additionally, PZ induces long range conformational changes in ZPI that brings in an added 2-fold enhancement of the rate of protease inhibition [14]. Therefore, PZ-ZPI binding interaction is also critical for the anticoagulant activity of ZPI. The binding hot spots between the two proteins were well described and shown to be centered on the sheet C, helix G and the C-terminus of helix A of ZPI [15]. Therefore, disrupting PZ-ZPI interface interactions can also be utilized as another efficacious way to suppress the activity of ZPI.

Molecular Dynamics (MD) simulation has been instrumental in revealing structural details of protein dynamics which are otherwise impossible to achieve by experimental methods. Thus, the over-reaching goal of the present work is to probe the conformational dynamics of RCL in ZPI employing all atom MD simulation to enable the prediction of residues critical for the inhibitory activity of ZPI-PZ and/ or ZPI. We performed MD simulation on the wild type (WT), Glu371 (E371K, E371Q and E371R, E371K-K318E) and Ser359 (S359Y, S359F, S359I) mutants of human ZPI. Our study revealed that the flexibility of RCL is connected to the overall stability of strand 5A that is contributed by both the Glu371 and Ser359 residues. The overall RCL dynamics of the E371 point variants did not change much in comparison to the WT due to compensatory interactions provided by the adjacent acidic residues, Glu368 and Asp370. However, altering the polarity of the electrostatic potential while retaining the stabilizing interactions of Glu371 through a double mutant K318E-E371K resulted in a significantly altered RCL dynamics. Highly flexible RCL was also observed in the Ser mutants, S359F and S359I, presumably due to loss of H-bond interactions made by the side chain of Ser359. S359Y, however, showed stability comparable to WT as the side chain of Tyr was able to maintain the stabilizing hydrogen bonding interaction. Next, using PCA analysis, we were able to observe that the stability of strand S5A not only affects RCL dynamics but also the PZ binding hotspots. Collectively, the current study provides a new perspective on the structural basis for efficient protease inhibition by ZPI.

## Materials & Methods

### System Preparation and MD simulation

The atomic coordinates of human wild-type ZPI were taken from the crystal structure by Wei *et al*., (PDB ID: 3F1S) [15] owing to its higher resolution. Atomic coordinates of the 4 variants E371K, E371Q, E371R and K318E-E371K were all obtained by *in silico* mutation of the wild type using Coot [16]. MD simulation was performed on the generated systems using AMBER 12 [17] with AMBER ff99 SB force field [18]. All the structures were energy minimized for 2000 steps using steepest descent and conjugate gradient algorithm. The energy minimized structures were then hydrated in a cubic water box comprising of TIP3P water molecules [19]. Charge neutralization was performed by adding requisite number of Cl^−^ ions. Electrostatic interactions were calculated with a cut-off distance of 12 Å using Particle mesh Ewald method [20]. Initial minimization steps were carried out to avoid bad contacts. This was followed by equilibration using NVT ensemble at 300 K for about 500 ps. The density of the systems was then equilibrated using NPT ensemble at 1 atm pressure for 1 ns. The time step used was 2 fs throughout the minimisation-equilibration-production run phase. After the energy values and density were seen to converge, the systems were subjected to 100 ns production using NPT ensemble at 300 K and 1 atm pressure. The coordinates were saved every 2 ps. The trajectories were visualized using VMD [21] and analyses such as RMSD, RMSF, hydrogen bond occupancy over time, inter atomic distances, secondary structure analysis were performed using CPPTRAJ [22] module of AMBER12.

### PCA analysis

In order to extract the principal components of the residue movements, principal component analysis was carried out on the dynamic trajectories of the wild type and mutants of ZPI using the ProDy package which is interfaced as the “normal mode wizard” [23] to the VMD 1.9.1 molecular graphics software. Next, the trajectories were aligned using the RMSD trajectory tool in VMD. Five hundred frames were considered in Principal component analysis (PCA) calculations. Only the 10 largest amplitude motions were calculated and porcupine plots for the first three modes were generated showing Eigen vectors along every fourth residue for clarity. The residue wise cross correlation map was also generated during the analysis for all the Cα atoms.

### Analysis of residue interaction networks (RINs)

The average structure derived from the 50 ns trajectory of each system was used to construct the RINs interactively using RING 2.0 webserver [24]. The RINs were generated based on a probe that detects all possible interactions such as interatomic contact, hydrogen bonds, salt bridges, pi–pi interaction, etc., between two atoms in contact. The generated nodes and edges were visualized using the RIN Analyzer plugin integrated with Cytoscape [25]. The network of interaction at the hinge region alone were extracted and presented.

All other visualization and analyses were carried out using Coot and all structural diagrams were created using PyMoL [26].

## Results and Discussion

MD simulation was started on the crystal structure of human ZPI (PDB ID: 3F1S) [15]. This structure was chosen owing to its higher resolution compared to the other reported structure of ZPI (PDB ID: 3H5C) [14]. In this structure, ZPI exists as a complex bound to PZ-GLA deleted mutant with an incomplete EGF1 domain. The structure of ZPI in the complex lacks the first 38 residues at the N-terminus but the typical serpin fold of ZPI can be clearly observed, with 3 main β-sheets and a fully exposed uncleaved RCL (Figure 1A). The RCL of ZPI is made up of residues 369–392 and the conformation of the P1 Tyrosine (that undergoes proteolytic cleavage during the conformational transition) present at position 387 is also clearly resolved. [15]. Glu371, a conserved hinge residue of ZPI, is located at the breach position which is at the junction between the top of β strand 5A and the base of the RCL. The corresponding Z variant (E342K) of antitrypsin (α1-AT) is the most common pathological variant that leads to α1-AT deficiency, liver disease and emphysema. The diseases caused by E342K mutant has been attributed mostly to the increased polymerization propensity due to an open and highly flexible conformation near the breach region. It was later found that the aberrant conformation of Z α1-AT is majorly due to loss of stabilizing interactions in strand S5A rather than to the change in the charge of the Glu342 residue [27]. The Ser 359 residue, on the other hand is equivalent to a highly conserved Ser 365 residue located at the N-terminal end of strand S5A in antithrombin. S365L variant of antithrombin was found to lack the inhibitory potential of the serpin and had a higher susceptibility to form covalent dimers in plasma through an unknown mechanism [28]. The simulations of the variants in ZPI were started by mutating Glu371 to Lys, Arg and a Gln residue and Ser359 **t**o Tyr, Phe and Ile using the original WT coordinates. For each system, independent simulations of 100 ns were performed. In all cases, the majority of secondary structure remained stable throughout the simulation. However, helix hF, the top of strands S6A and S5A, a 3_10_ helix connecting sheet C and sheet A, the turn connecting strands 3C and 4C, as well as the RCL showed a high degree of flexibility (Figure 1B). The root mean square deviations (rmsd) from the starting geometries as a function of simulation time indicate that all the structures converge and stabilize after 50 ns (Figure 2). Therefore, all our analyses were performed using the 50-100 ns trajectories.

**Figure 2.**
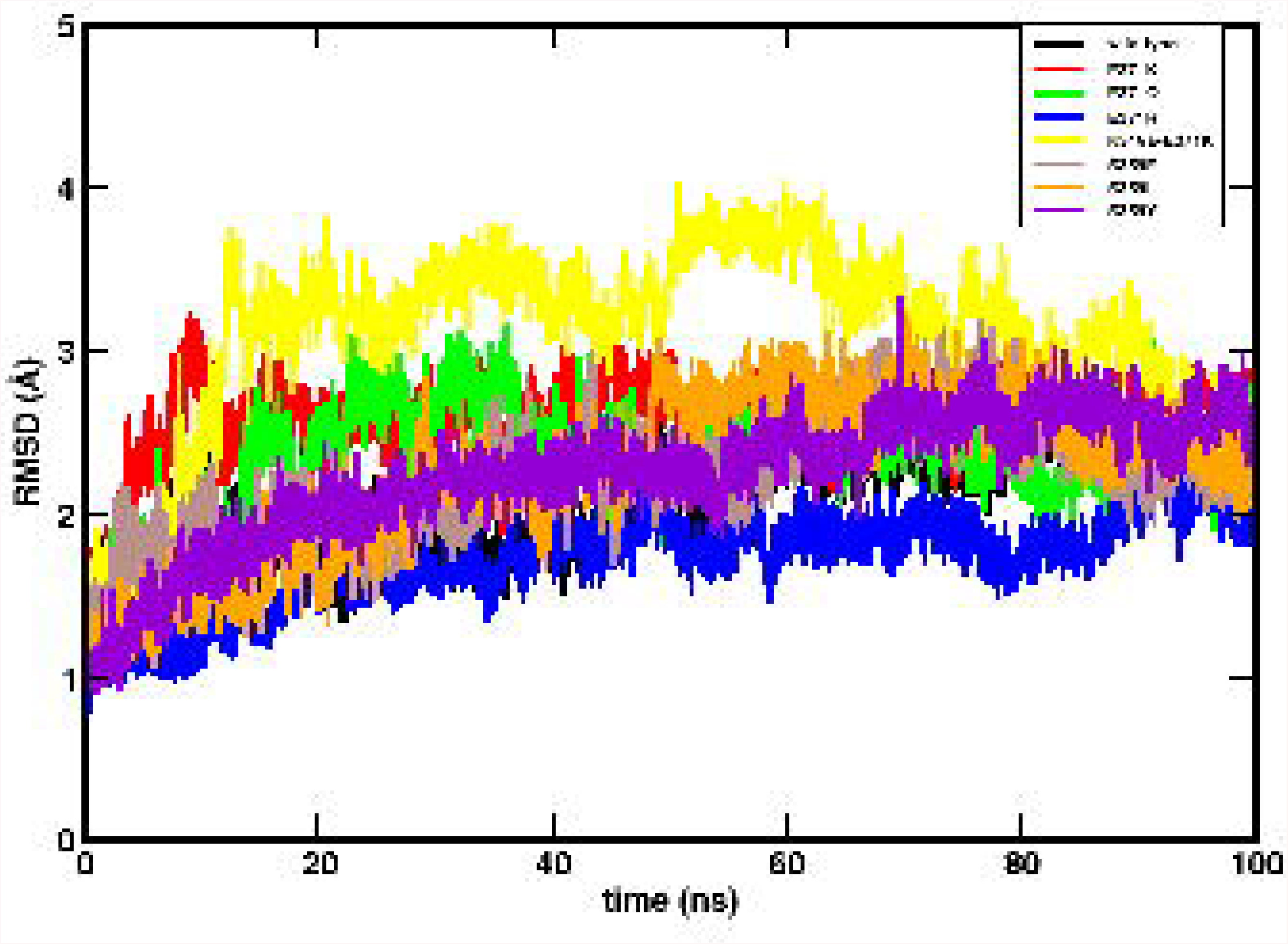
The time evolution for the root mean square deviation (RMSD) of the simulated structures with respect to the first frame of simulation has been shown for the different systems.

### Comparison of Interactions at the breach region

Throughout the simulation with the WT, the hydrogen bonding interactions between strands S3A, S5A were maintained. Likewise, the electrostatic interactions of Asp370 (S5A) with Arg372 (RCL) and Lys220 (S3A), and that of Glu371 (RCL) with Lys318 (S6A) were also retained. The simulation data for the fully optimized E371K mutant showed slight strand separation by ≥ ~ 1.5 Å between S3A and S5A in comparison to that of the final 100 ns simulated structure of WT (Figure S1). This result is quite similar to that of α-1 AT and its Z variant [11]. Changes in this region affected the rate of strand insertion post RCL cleavage in the Z-variant. However unlike in α-1 AT, no significant changes in the hydrogen bonding pattern between the strands were observed in ZPI (Figure 3). Consistently, the expected interaction between the conserved Trp221 and the backbone amide of Asp370 is retained in both the WT and the E371K variant throughout the simulation (Figure 3). The major difference in E371K compared to the WT, is the loss of the interaction between Glu371 of the RCL and Lys318 of strand S6A. However, the loss of this interaction is rescued by the interaction of the adjacent RCL residue Asp370 with Lys318. Additionally, the E371K residue makes a new intra-RCL salt bridge interaction with Asp370, whereas Arg372 residue of the RCL now inserts into the breach region and makes interactions with Glu368 of strand S5A (Figures 3 and 4). Consequently, the proximal hinge of the RCL appears to be partially inserted into the top of the β-sheet between strands S3A and S5A. This interaction may be the cause for the observed strand separation in E371K (Figure 3 and Figure S1).

**Figure 3.**
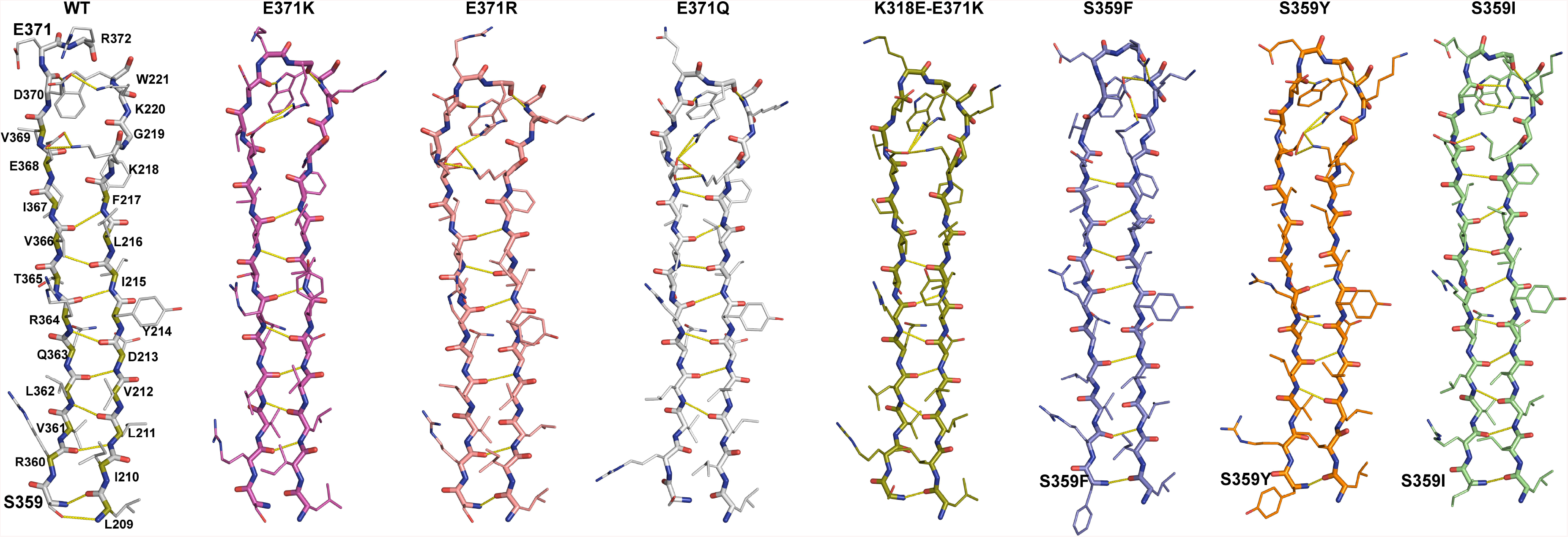
Inter-strand H-bonds between strands S3A and S5A in the WT and the mutants. Inter-strand H-bonds between strands s3A and s5A of representative time-averaged structure (over the last 50 ns of simulation) of WT and different E371 and S359 variants. No significant alteration in the hydrogen bonding pattern between the strands were observed in the WT and the variants. The conserved Trp221 can be found to interact with the backbone amide of Asp370 throughout the simulation in all the proteins.

**Figure 4.**
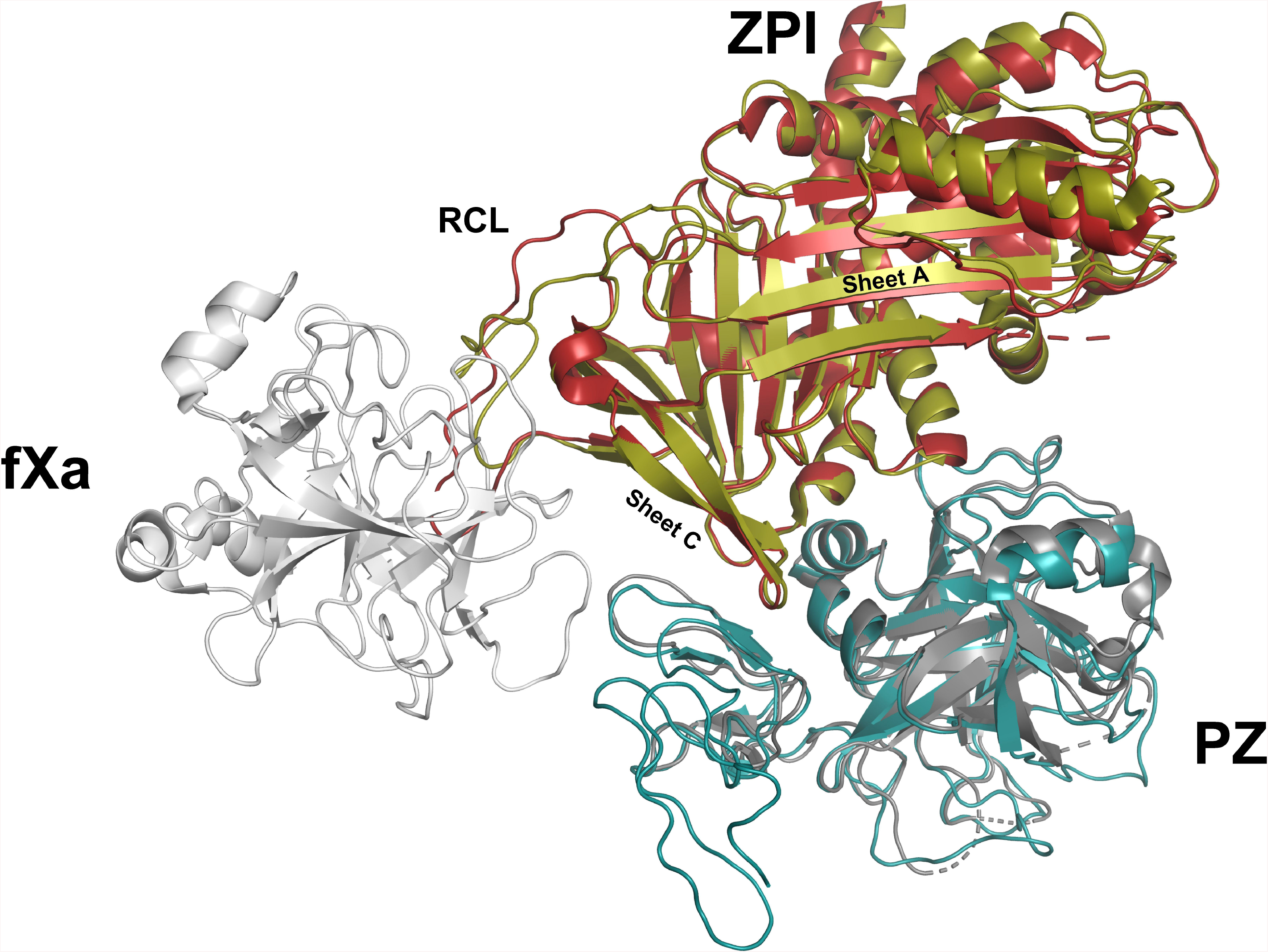
Cartoon showing a fXa-ZPI-PZ model. Two different PZ-ZPI complex structures are shown as superposed ribbons. The ZPI-PZ complex lacking the EGF1 domain is shown in maroon and grey, respectively (PDB ID: 3F1S), and the complex with the EGF1 domain is shown in green and blue, respectively (PDB ID: 3H5C). In the ternary complex model with the ZPI-PZ complex lacking the full EGF1 domain, the RCL of ZPI (maroon) does not appear to be complementary to the fXa binding region.

In this context, it is to be noted that the Arg372 residue was found to be exposed to the solvent in the two different crystal structures of WT ZPI. In the other reported structure of the WT (PDB ID: 3H5C) the RCL exists in an extended conformation and the solvent exposed Arg372 residue makes a hydrogen bond interaction with the main chain carbonyl of the RCL residue Glu375 [14]. Since the two crystal forms captured the RCL in two different conformations, we decided to test for the most compatible RCl conformation for effective docking onto fXa. For that, we made a model of the ternary Michaelis complex of fXa with ZPI and PZ, similar to the model proposed by Huang et al., based on the AT-factor Xa ternary complex (Figure 4) [14]. It appears that interaction between the RCL of ZPI and fXa would be more favorable when the RCL assumes an extended conformation where the Arg372 makes the intra-RCL hydrogen bond (PDB ID: 3H5C). In such a scenario, it is likely that the solvent exposed conformation of Arg372 is important to facilitate formation of the required active/extended conformation of the RCL for proper docking onto fXa. Notably, Arg372 is a completely conserved residue in ZPI and is unique only to this family of serpins suggesting that the conformation of this residue might be functionally important in regulating ZPI activity (Figure 5).

**Figure 5.**
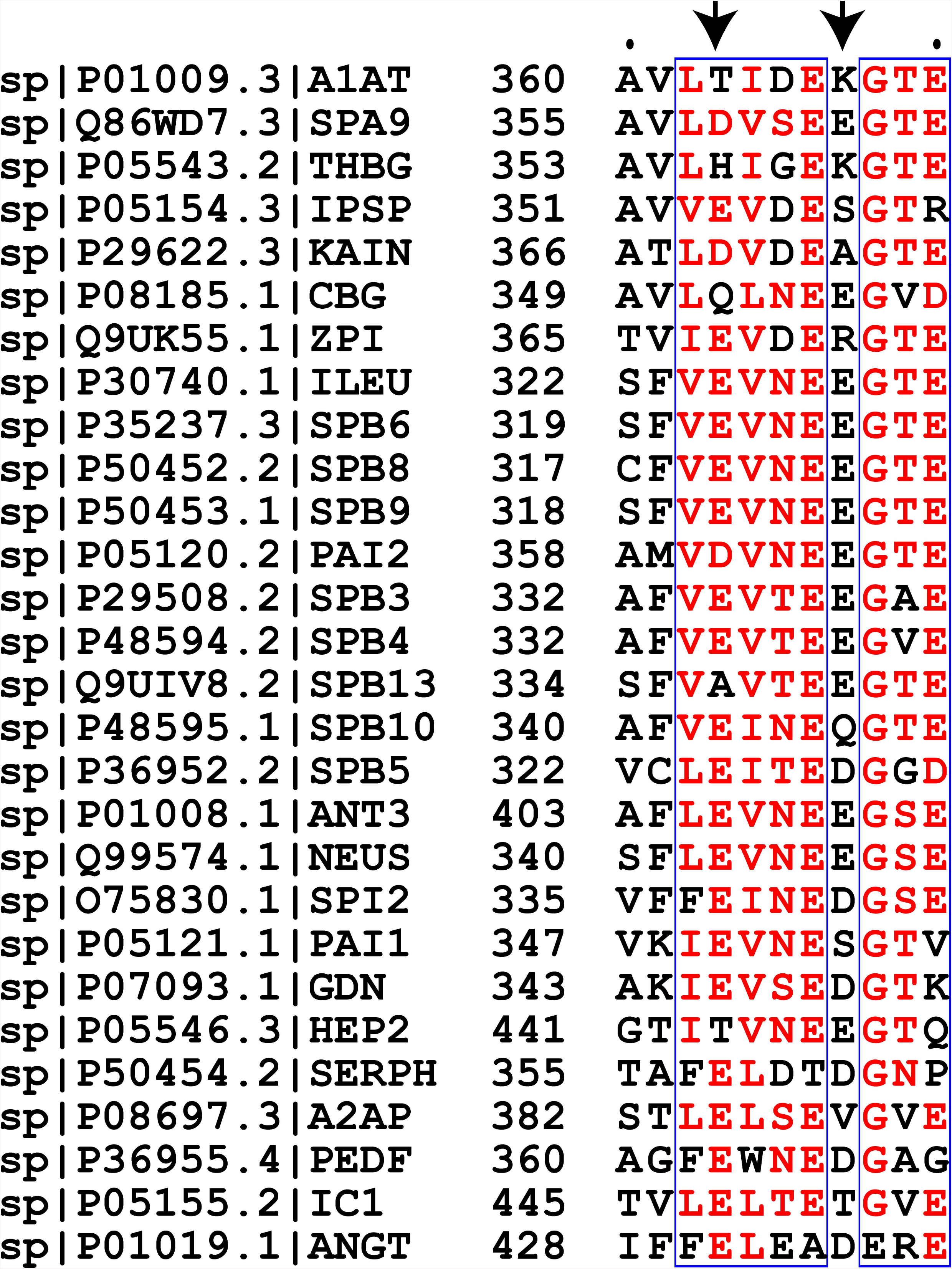
Multiple sequence alignment of different humans SERPINS shows that the interaction between Arg372 and Glu368 is unique to ZPI.

### Effect of the E371 mutants on RCL flexibility

Changes in the breach region such as loosening up of the inter-strand interactions in the Z-variant of α-1 AT displayed higher RCL flexibility and was proposed to contribute to its higher rate of polymerization, since loop insertion would be easier in the slightly open state when the breach region is solvent exposed. We expected a similar effect on the RCL of the E371K mutant of ZPI. Surprisingly, the flexibility of the RCL in the E371K variant was comparable to the WT ZPI as shown in figure (Figure 6). This indicates that introducing a positive charge at Glu371 despite the energetically unfavorable proximity to Lys318 or Arg372 does not significantly affect the RCL dynamics of ZPI in contrast to the Z variant of α-1 AT. To test the observation further, we altered the physicochemical properties of the side chain at position 371 by generating the E371R and E371Q mutants modeled onto ZPI and performed MD studies on those variants. The substitutions are stereo chemically conservative but alter only the strength of the electrostatic potential at this position. Both E371Q and E371R were found to be stable after 50 ns of simulation (Figure 2). Once again the stability of the RCL was found to be stable in these two mutants and comparable to the WT protein (Figure 6). In the E371Q variant, the glutamine residue makes a H bond interaction with the Lys318 of strands S6, whereas in the E371R variant, arginine forms a favorable stable stacking interaction with a neighboring Pro227 residue of strand 4C (Figure S3). Similar to E371K, we found that Arg372 in both E371Q and E371R moves away from the variants and inserts into the top of the sheet and makes salt bridge interactions with Glu368 (Figures 3, 7 and S2). It appears that the compensatory interactions made by Asp370, the E371variants, and Arg372 at the proximal end of the RCL with strands S6A and S5A rescues the stability of the RCL in the E371 variants of ZPI. However, when we performed a hydrogen bond interaction analysis of the RCL with the last 5 frames, we noticed that the loop had a greater propensity to form intra-loop H bond in the mutants in comparison to the WT (30% in the WT, 40% in E371Q, 45% in E371K, and 60% in the E371R variant). This suggests that in the mutants, specifically E371R, the energetics associated with altering the RCL conformation for appropriate docking onto fXa would be altered. Consequently, it is likely that the RCL might not be able to undergo the necessary changes required for efficient interaction with fXa and therefore the rate of spontaneous loop insertion could be affected.

**Figure 6.**
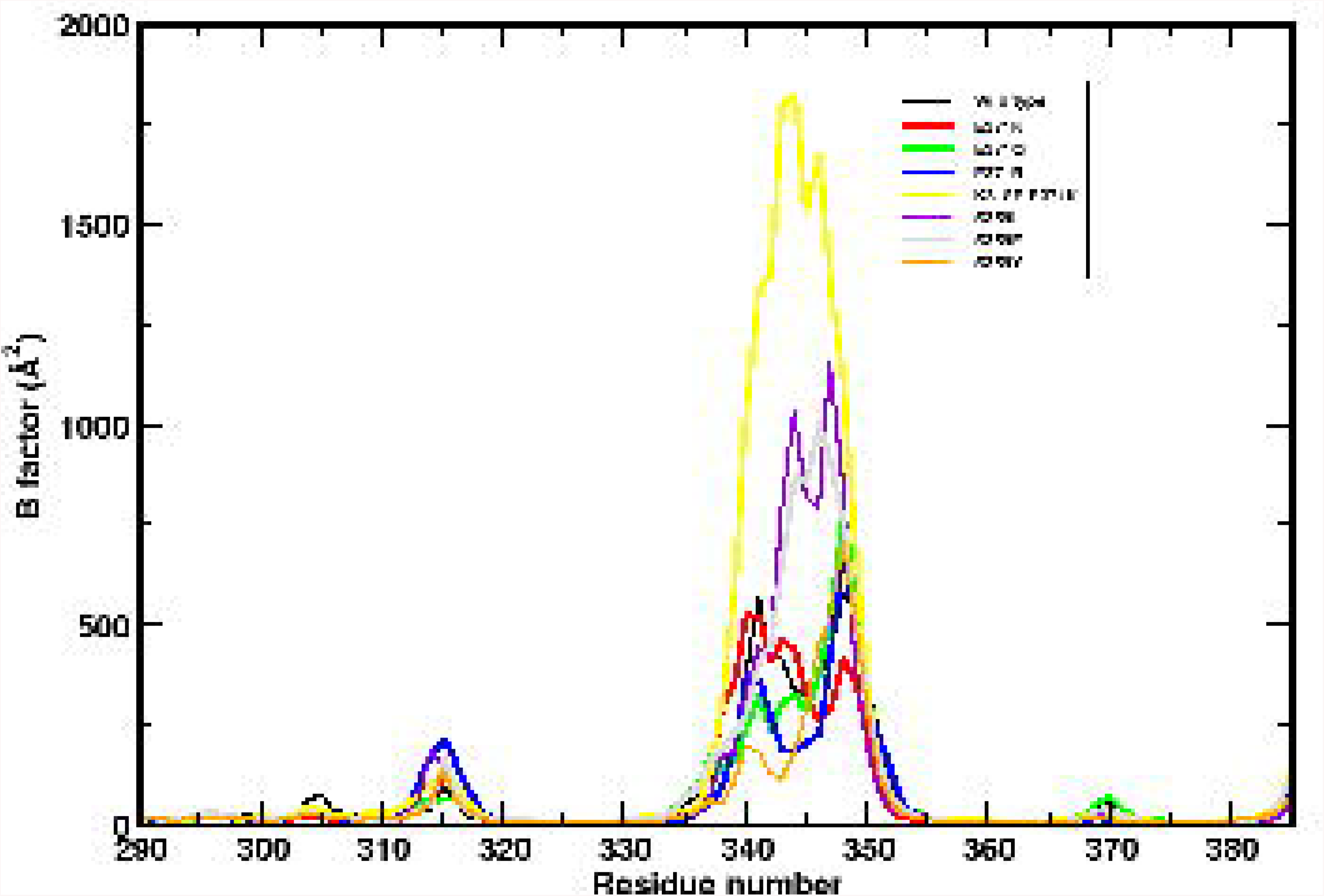
Backbone RMSF of the RCL residues in the WT and the mutants.

**Figure 7.**
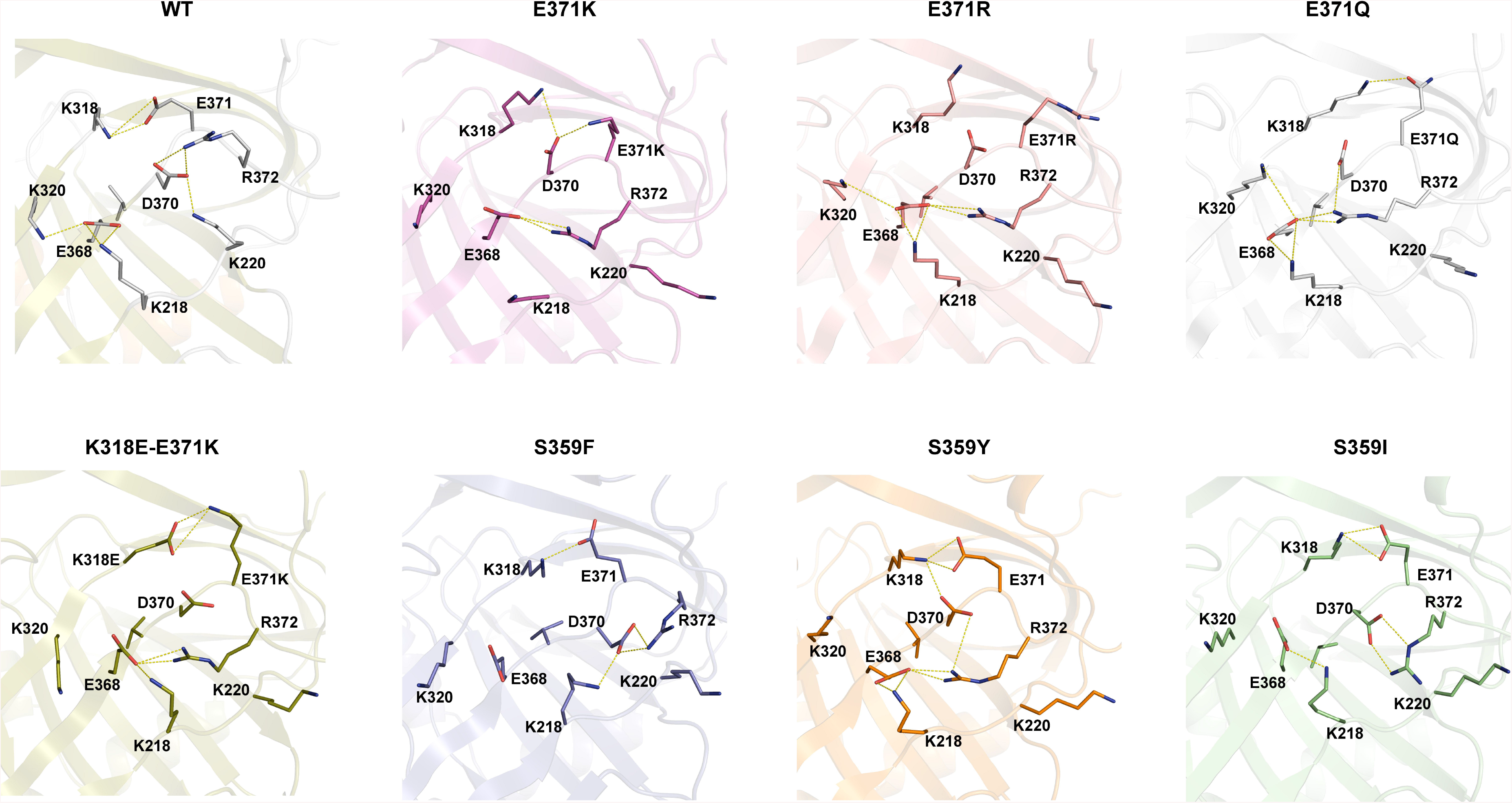
Electrostatic interactions at the breach region. Representative conformations of residues at the breach region from time-averaged structures of the last 50 ns of the simulation. (A) WT, (B) E371K, (C) E371R, (D) E371Q, (E) K318E-E371K, (F) S359F, (G) S359Y and (H) S359I. In the Glu371 variants, the RCL appears to be partially inserted into the top of the β-sheet between strands S3A and S5A due to the interaction of Arg372 with Glu368 of strand S5A. Also, the loss of the salt bridge between Glu371 and Lys318 is rescued by the interaction of the adjacent RCL residue Asp370 with Lys318 in the variants.

In an alternate experiment, we generated the double mutant K318E-E371K. In this case, we wanted to test the effect of switching the electrostatic potential at the top of the breach while maintaining the net electrostatic potential to remain identical to the WT protein. This mutant showed identical strand separation to that of the E371K mutant (Figure S1). As expected, K318E and E371K made a salt bridge interaction like in the WT, and Arg372 inserted into the breach region to make the salt bridge interaction with Glu368. Since the overall interactions appeared similar to the WT, we expected the RCL dynamics to also resemble that of the WT. Unexpectedly, the RCL here displayed very high flexibility (Figure 6). The only major difference in this mutant is the unfavorable proximity of the K318E to Asp370 of the RCL and Glu368 of strand S5A. This perhaps contributes to a decreased overall stability of the strand S5A that in turn contributes to the increased flexibility of RCL. A closer observation of the surface electrostatic potential of the breach region clearly indicates that the region is highly negatively charged and it appears that introducing more negative behavior at position 318 adjacent to the already negatively charged breach region significantly affects RCL stability whereas introducing the positive charge at 371 is compensatory and does not significantly alter the stability of the RCL. Clearly, the effect of the analogous Z-variant of α-1 AT (E342K) will have a different effect on the functional dynamics of ZPI in comparison to α-1 AT or other homologous serpins.

### Disruption of interactions at the N-terminal end of strand S5A also affects RCL stability

Since we were able to observe a correlation between the stability of strand S5A and the RCL of ZPI, we next attempted to alter the stability of strand S5A without interfering the interactions at the breach region. Apart from the interactions made at the breach and the shutter, Ser359 is the only other residue that makes a side chain mediated interaction at the N-terminal end of S5A. Ser359 is a highly conserved residue and mutations in this residue in antithrombin result in the formation of unstable complexes with fXa [28]. In an attempt to destabilize the interaction made by Ser359, we mutated the residue to Tyr, Phe and Ile and carried out the MD simulations. When we analyzed the RCL flexibility, it could be observed that the S359I mutant significantly affected the RCL flexibility followed by the S359F variant. The S359Y variant had comparable RCL flexibility to that of the WT (Figure 6). The observation that the two hydrophobic residues affected RCL flexibility suggests that the hydrogen bond interaction made by the hydroxyl group of the Ser (WT) and that of the Tyr (S359Y variant) is important for the S5A stability. Our findings imply that the interactions at the distal end of strand S5A is also crucial for RCL flexibility and any slight perturbations on the interactions made by strand S5A can affect the rate of RCL insertion for effective protease inactivation.

### Principal component analysis reveals that the PZ binding hotspot and the RCL undergo correlated motions

ZPI is a unique serpin in that it gets activated by PZ whilst most other serpins of hemostatic system get activated by heparin [29,30]. However, heparin was also reported to act modestly as a cofactor of ZPI for both fXa and fXIa inhibition [31]. Interestingly, PZ binds to ZPI on the face closest to strand S6A and opposite to the heparin binding site [32,33]. Since changes in Lys318 present in S6A affected RCL dynamics, we attempted to see if the mutations under study, present at the breach region might affect the PZ binding hotspots of ZPI.

Several ionic interactions contribute to the binding interface of PZ and ZPI, but the most significant contribution comes from the packing of the hydrophobic residues Tyr240 and Met71 of ZPI into the complimentary cavities in PZ (Figures 7 and 8) [34]. The hotspot residue closest to the breach region is Tyr240, present at a turn between strands 3C and 4C (Figure 8). A closer observation of the interactions made by the residues Glu371 and Lys318 reveals that, apart from the salt bridge interaction with Lys318, the side chain of Glu371 makes an H bond interaction with the side chain of Thr230 of strand 3C. Lys318 also makes stacking interactions with Pro246 of strand 4C. Thr230 can also interact with the RCL through the 3_10_ helix adjacent to it (Figure 8). Thus, it is likely that changes in the network of interactions in this region can be relayed to the PZ-binding hotspot residue Tyr240, the RCL and the sheet A through the breach region. Therefore, we decided to employ principal component analysis (PCA) to test if the motions of these three regions are correlated in the WT and the K318E-E371K variant where the RCL displays highest flexibility. Interestingly, Z mutation of α1-AT was also reported to alter the dynamics of regions distant to the mutation site [35].

**Figure 8.**
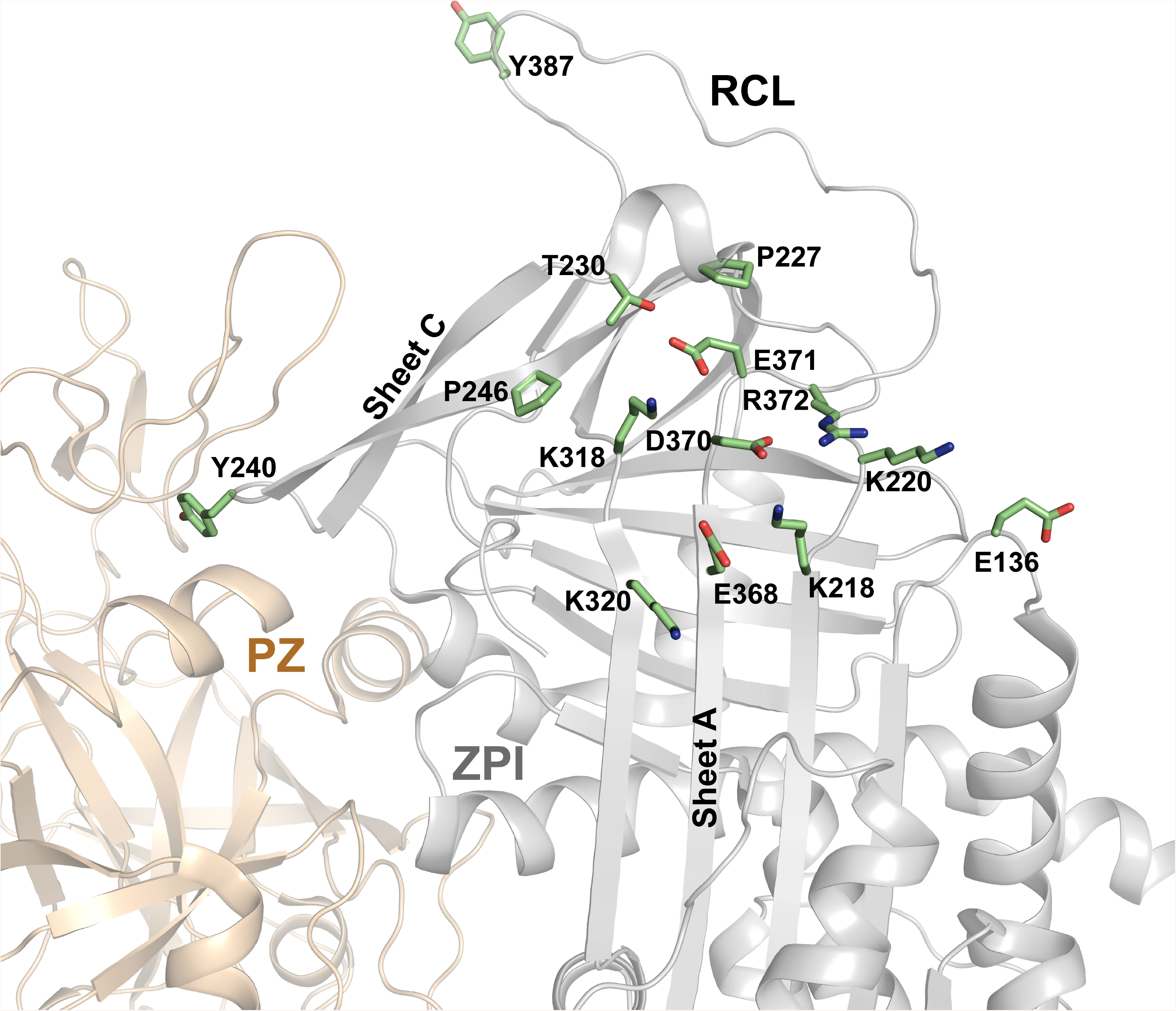
Close-up view of the PZ-ZPI interface near the hinge region. PZ (cream) and ZPI (grey) are shown as cartoons. Tyr240 of sheet C of ZPI can be seen interacting with PZ. Residues present in sheet A at the hinge region and the P1 Tyr387 of ZPI are represented as green sticks (PDB ID: 3F1S).

Three low frequency modes from the PCA of the structures were generated (Figure 9). The protein motions along specific directions in each of these modes is given by eigenvectors in the porcupine plot. A clear difference in the directions of motion can be seen in the comparative motion in both the proteins perhaps as a result of change in the network of interactions in the breach region (Figure 9A). The residue based mobility plots across three different modes is shown in figure 9B and the list of residues that contributed maximally towards the motions across each of these modes is shown in Table 1. In the WT, a clear correlation in the motion of the RCL and the PZ binding Tyr240 region could be observed in mode1 itself whereas in the K318E-E371K mutant, the RCL showed complete lack of correlated movements (Figure 9B). However, when we observe the residue wise cross correlation plot, we can see that while these two regions show slight negative correlation in the WT, there are extensive correlation in both directions in the double mutant (Figure 9C). In order to resolve this ambiguity, we generated the residue wise interaction network (RIN) for the WT and the mutant. The changes in the interaction between the residue at 371 position and the other regions connecting to sheet A and sheet C could be more obviously detected by observing the RIN plot (Figure 9D). Clearly, the network observed in the WT is disrupted in the mutant. This alteration in the networks most likely contributes to the observed changes in fluctuation captured by the 3 principal component modes. There are a total of 10 nodes and 12 edges that include the first and second neighbors interacting with E371 in the WT, whereas it is 18 nodes and 25 edges for the double mutant K318E-E371K. The increased connection of K318E and E371K with several regions in the double mutant perhaps results in the flexibility of the RCL being sensitive to even very slight changes in the protein that are not captured in the mobility plot. Thus the findings here indicate that the regulation of correlated motion is lost in the double mutant in comparison to the WT.

**Figure 9.**
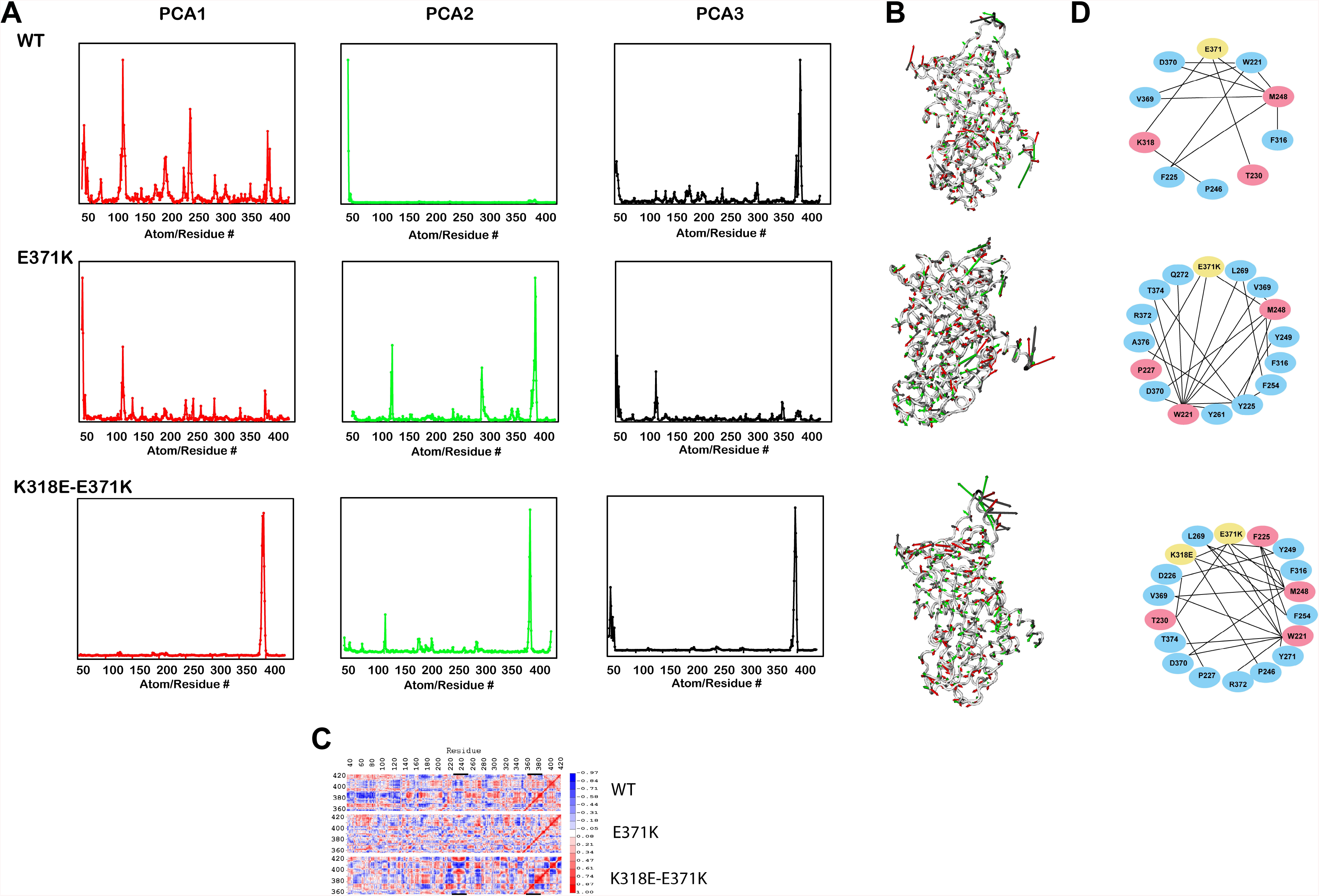
A) Residue based mobility plot showing mobility of different residues across the respective modes. B) Porcupine plots showing prominent motions along the three PCA modes. Eigen vectors showing the direction of prominent motions across modes 1, 2 and 3 are shown as red, green and black arrows, respectively. C) Residue wise cross correlation with the hinge and RCL regions in the WT and mutants. D) Comparison of residue interaction networks between the wild and the variants of ZPI highlighting changes in network interaction due to the mutation at the hinge region. The E371 residue and its variants are highlighted in yellow. The first and second neighbors are shown in pink and blue respectively.

**Table 1.0.**
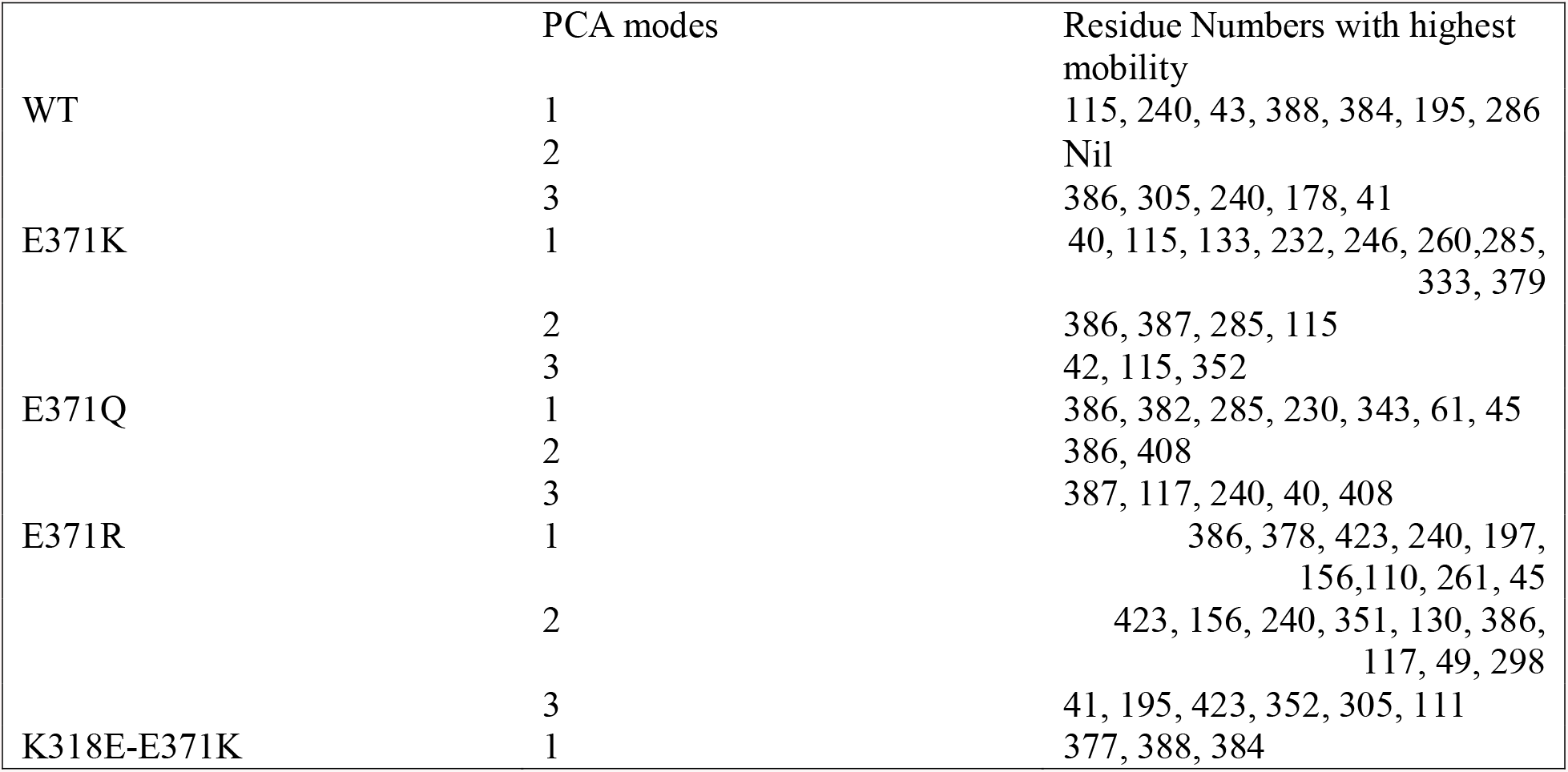
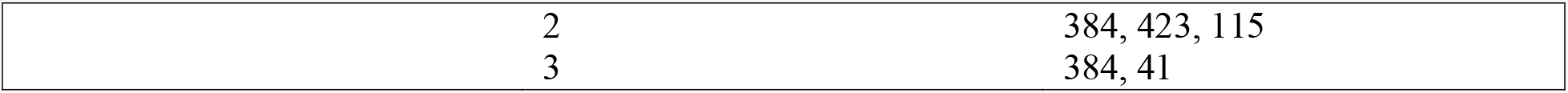
Residues that contribute to maximum motions in the WT and variants of ZPI in the first three principal modes.

|  | PCA modes | Residue Numbers with highest mobility |
| --- | --- | --- |
| WT | 1 | 115, 240, 43, 388, 384, 195, 286 |
|  | 2 | Nil |
|  | 3 | 386, 305, 240, 178, 41 |
| E371K | 1 | 40, 115, 133, 232, 246, 260, 285, 333, 379 |
|  | 2 | 386, 387, 285, 115 |
|  | 3 | 42, 115, 352 |
| E371Q | 1 | 386, 382, 285, 230, 343, 61, 45 |
|  | 2 | 386, 408 |
|  | 3 | 387, 117, 240, 40, 408 |
| E371R | 1 | 386, 378, 423, 240, 197, 156, 110, 261, 45 |
|  | 2 | 423, 156, 240, 351, 130, 386, 117, 49, 298 |
|  | 3 | 41, 195, 423, 352, 305, 111 |
| K318E-E371K | 1 | 377, 388, 384 |

|  |  |
| --- | --- |
| 2 | 384, 423, 115 |
| 3 | 384, 41 |

## Conclusion

ZPI metastability is an important factor that dictates its function. Specifically, RCL flexibility and loop insertion to form a part of the beta sheet A is critical for protease interaction and its corresponding inactivation. While extensive mutagenesis studies have identified residues important for successful loop insertion in α1-antitrypsin and antithrombin, similar studies are lacking for ZPI. For example, the proximal hinge Glu residue on the RCL in α1-AT was proposed to be important for maintaining loop flexibility and insertion and the Z variant of that residue (E342K) is implicated in many diseases. In another instance, the Ser365 present at the distal end of antithrombin sheet A was reported to be crucial for the rate of RCL insertion and mutation of the residue (S365L) was identified in patients with type II antithrombin deficiency. The corresponding Glu371and Ser359 are also conserved in ZPI. We thus decided to investigate the effect of substituting E371 and S359 on the flexibility of the RCL in order to identify if the residues can also play a similar role in ZPI. Although the exact mechanism of loop insertion in ZPI is not known, we saw two different RCL conformations in two different crystal structures of ZPI in complex with PZ. Since two different RCL conformations of ZPI in two different crystal structures were observed in the PZ bound state, it is quite likely that PZ also plays some role in maintaining proper conformational dynamics of ZPI RCL. However, the residues required to relay the signal of PZ binding and activation for fXa inhibition is not known. Therefore, we also probed if the mutation in the breach region can play a role in relaying PZ binding to the RCL of ZPI.

Our study demonstrated that unlike the Z variant of α1-AT, there is no appreciable change in the RCL as well as overall stability of E371K variant compared to the WT. The contrasting charge distribution near the breach region of ZPI (having mostly negatively charged residues) compared to α1-AT (having mostly positively charged residue) was considered to be the prime reason for such disparity (Figure S4). Hence, it is likely that the E371K variant of ZPI would behave similarly like the WT protein. However, the double mutant (E371K-K318E) indicated that switching the electrostatic potential at the breach region resulted in a variant with very high RCL flexibility. Thus, although Glu371 interactions are indeed required for stabilizing the native conformation of ZPI, the basicity of Lys318 appears to be most important for this purpose. This was further corroborated by PCA analysis which showed a complete lack of regulation of correlated motion between the RCL and the rest of the protein in the double mutant compared to the WT, whereas correlated motion is completely retained in E371K. Another important residue making stabilizing interactions at strand S5A was shown to be S359 as mutation of Ser to hydrophobic residues such as Phe and Ile resulted in enhanced RCL flexibility. Notably, mutation of the homologous residue in antithrombin would result in a variant, S365L, which was reported to get precipitated as inactive dimers. Therefore, perturbations of stabilizing interactions made by Lys318 and Ser359 would affect the stability of strand S5A which in turn would result in a highly flexible RCL of ZPI with a potential for higher rate of polymerization and lack of regulation of its activity.

## Acknowledgement

Authors wish to thank Dr Rajeswari Appadurai, IISC Bangalore, for her helpful comments and suggestions. The authors also acknowledge the computing facility of the P. G. Senapathy Centre, IIT Madras.

## References

1. Xin Han, Ryan Fiehler, George J. Broze J (2000) Characterization of the protein Z– dependent protease inhibitor. Blood 96:3049–3055

2. Gettins PGW (2002) Serpin Structure, Mechanism, and Function. Chem Rev 102:4751–4803

3. Johnson DJ, Huntington JA (2004) The influence of hinge region residue Glu-381 on antithrombin allostery and metastability. J Biol Chem 279 (6):4913–4921. doi:10.1074/jbc.M311644200

4. Huntington JA (2011) Serpin structure, function and dysfunction. J Thromb Haemost 9:26–34

5. Huang X (2019) Engineering a protein Z-dependent protease inhibitor (ZPI) mutant as a novel antagonist of ZPI anticoagulant function for hemophilia treatment. J Thromb Haemost 17 (10):1655–1660. doi:10.1111/jth.14610

6. Girard TJ, Lasky NM, Grunz K, Broze GJ, Jr. (2019) Suppressing protein Z-dependent inhibition of factor Xa improves coagulation in hemophilia A. J Thromb Haemost 17 (1):149–156. doi:10.1111/jth.14337

7. Polderdijk SG, Adams TE, Ivanciu L, Camire RM, Baglin TP, Huntington JA (2017) Design and characterization of an APC-specific serpin for the treatment of hemophilia. Blood 129 (1):105–113. doi:10.1182/blood-2016-05-718635

8. Julie A Peterson SAM, Alan E Mast (2016) Targeting TFPI for hemophilia treatment. Thromb Res 141:S28–30

9. Raja Prince LB, Mirko Manetti, Daniela Melchiorre, Irene Rosa, Natacha Dewarrat, Silvia Suardi, Poorya Amini, Joś e A. Ferń andez, Laurent Burnier, Claudia Quarroz, Maria Desiŕe Reina Caro, Yasuhiro Matsumura, Johanna A. Kremer Hovinga, John H. Griffin, Hans-Uwe Simon, Lidia Ibba-Manneschi, François Saller, Sara Calzavarini, Anne Angelillo-Scherrer (2018) Targeting anticoagulant protein S to improve hemostasisin hemophilia. Blood 131 (12):1360–1371

10. K. John Pasi SR, Pencho Georgiev, Tim Mant, Michael D. Creagh, Toshko Lissitchkov, David Bevan, Steve Austin, Charles R. Hay, Inga Hegemann, Rashid Kazmi, Pratima Chowdary (2017) Targeting of Antithrombin in Hemophilia A or B with RNAi Therapy. N Engl J Med 377:819–828

11. Kass I, Knaupp AS, Bottomley SP, Buckle AM (2012) Conformational properties of the disease-causing Z variant of alpha1-antitrypsin revealed by theory and experiment. Biophys J 102 (12):2856–2865. doi:10.1016/j.bpj.2012.05.023

12. Tanusree Sengupta NM (2016) Phosphatidylserine and Phosphatidylethanolamine Bind to Protein Z Cooperatively and with Equal Affinity PLoS One 11

13. Huang X, Swanson R, Kroh HK, Bock PE (2019) Protein Z-dependent protease inhibitor (ZPI) is a physiologically significant inhibitor of prothrombinase function. J Biol Chem 294 (19):7644–7657. doi:10.1074/jbc.RA118.006787

14. Huang X, Dementiev A, Olson ST, Gettins PG (2010) Basis for the specificity and activation of the serpin protein Z-dependent proteinase inhibitor (ZPI) as an inhibitor of membrane-associated factor Xa. J Biol Chem 285 (26):20399–20409. doi:10.1074/jbc.M110.112748

15. Zhenquan Wei, Yahui Yan, Robin W. Carrell, Zhou A (2009) Crystal structure of protein Z–dependent inhibitor complex shows how protein Z functions as a cofactor in the membrane inhibition of factor X. Blood 114 (17):3662–3667

16. P. Emsley, B. Lohkamp, W. G. Scottc, Cowtan K (2010) Features and Development of Coot. Acta Crysta D66:486–501

17. D.A. Case TAD, T.E. Cheatham, III, C.L. Simmerling, J. Wang, R.E. Duke, R. Luo, R.C. Walker, W. Zhang, K.M. Merz, B. Roberts, S. Hayik, A. Roitberg, G. Seabra,J. Swails, A.W. Götz, I. Kolossváry, K.F. Wong, F. Paesani, J. Vanicek, R.M. Wolf, J. Liu,X. Wu, S.R. Brozell, T. Steinbrecher, H. Gohlke, Q. Cai, X. Ye, J. Wang, M.-J. Hsieh, G. Cui, D.R. Roe, D.H. Mathews, M.G. Seetin, R. Salomon-Ferrer, C. Sagui, V. Babin, T. Luchko, S. Gusarov, A. Kovalenko, and P.A. Kollma (2012) AMBER 12, University of California, San Francisco.

18. Wang J, Wolf RM, Caldwell JW, Kollman PA, Case DA (2004) Development and testing of a general amber force field. J Comput Chem 25:1157–1174

19. William L. Jorgensen, Jayaraman Chandrasekhar, Madura JD (1983) Comparison of simple potential functions for simulating liquid water J Chem Phys 79:926–935

20. Ulrich Essmann, Lalith Perera, Max L. Berkowitz, Tom Darden, Hsing Lee, Pedersen LG (1995) A smooth particle mesh Ewald method. J Chem Phys 103:8577–8593

21. Humphrey W, Dalke A, Schulten K (1996) VMD - Visual Molecular Dynamics. J Molec Graphics 14:33–38

22. Roe DR, Cheatham TE (2013) PTRAJ and CPPTRAJ: Software for Processing and Analysis of Molecular Dynamics Trajectory Data. J Chem Theory Comput 9:3084–3095

23. Bakan A, Meireles LM I B (2011) ProDy: Protein Dynamics Inferred from Theory and Experiments. Bioinformatics 27 (11):1575–1577

24. Piovesan D. MG, Tosatto S.C.E. (2016) The RING 2.0 web server for high quality residue interaction networks. Nucleic Acids Research 44 (W1): W367–374

25. Nadezhda T Doncheva KK, Francisco S Domingues, Mario Albrecht (2011) Analyzing and visualizing residue networks of protein structures. Trends Biochem Sci 36:179–182

26. The PyMOL Molecular Graphics System, Version 2.0 Schrödinger, LLC.

27. Huang X ZY, Zhang F, Wei Z, Wang Y, Carrell RW, Read RJ, Chen G-Q, Zhou A (2016) Molecular mechanism of Z α1-antitrypsin deficiency. J Biol Chem 291:15674–15686

28. Sonia Aguila GI, Irene Martínez-Martínez, Vicente Vicente, Steven T. Olson, Corral aJ (2017) Disease-causing mutations in the serpin antithrombin reveal a key domain critical for inhibiting protease activities. J Biol Chem 292 (40):16513–16520

29. Robert N. Pike AMB, Bernard F. le Bonnie, Frank C. Church (2005) Control of the coagulation system by serpins Getting by with a little help from glycosaminoglycans. FEBS J 272:4842–4851

30. Whisstock JC, Pike RN, Jin L, Skinner R, Pei XY, Carrell RW, Lesk AM (2000) Conformational changes in serpins: II. The mechanism of activation of antithrombin by heparin. J Mol Biol 301 (5):1287–1305. doi:10.1006/jmbi.2000.3982

31. Xin Huang ARR, George J. Broze, Jr., Steven T. Olson (2011) Heparin Is a Major Activator of the Anticoagulant Serpin,Protein Z-dependent Protease Inhibitor. J Biol Chem 286:8740–8751

32. Huang X, Zhou J, Zhou A, Olson ST (2015) Thermodynamic and kinetic characterization of the protein Z-dependent protease inhibitor (ZPI)-protein Z interaction reveals an unexpected role for ZPI Lys-239. J Biol Chem 290 (15):9906–9918. doi:10.1074/jbc.M114.633479

33. Yang L, Ding Q, Huang X, Olson ST, Rezaie AR (2012) Characterization of the heparin-binding site of the protein z-dependent protease inhibitor. Biochemistry 51 (19):4078–4085. doi:10.1021/bi300353c

34. Huang X, Yan Y, Tu Y, Gatti J, Broze GJ, Jr., Zhou A, Olson ST (2012) Structural basis for catalytic activation of protein Z-dependent protease inhibitor (ZPI) by protein Z. Blood 120 (8):1726–1733. doi:10.1182/blood-2012-03-419598

35. Hughes VA, Meklemburg R, Bottomley SP, Wintrode PL (2014) The Z mutation alters the global structural dynamics of alpha1-antitrypsin. PLoS One 9 (9):e102617. doi:10.1371/journal.pone.0102617

