## Supplemental FigureS1-S4 for "Effect of Variations in the Conserved Residues E371 and S359 on the Structural Dynamics of Protein Z Dependent Protease Inhibitor (ZPI): A Molecular Dynamic Simulation Study"

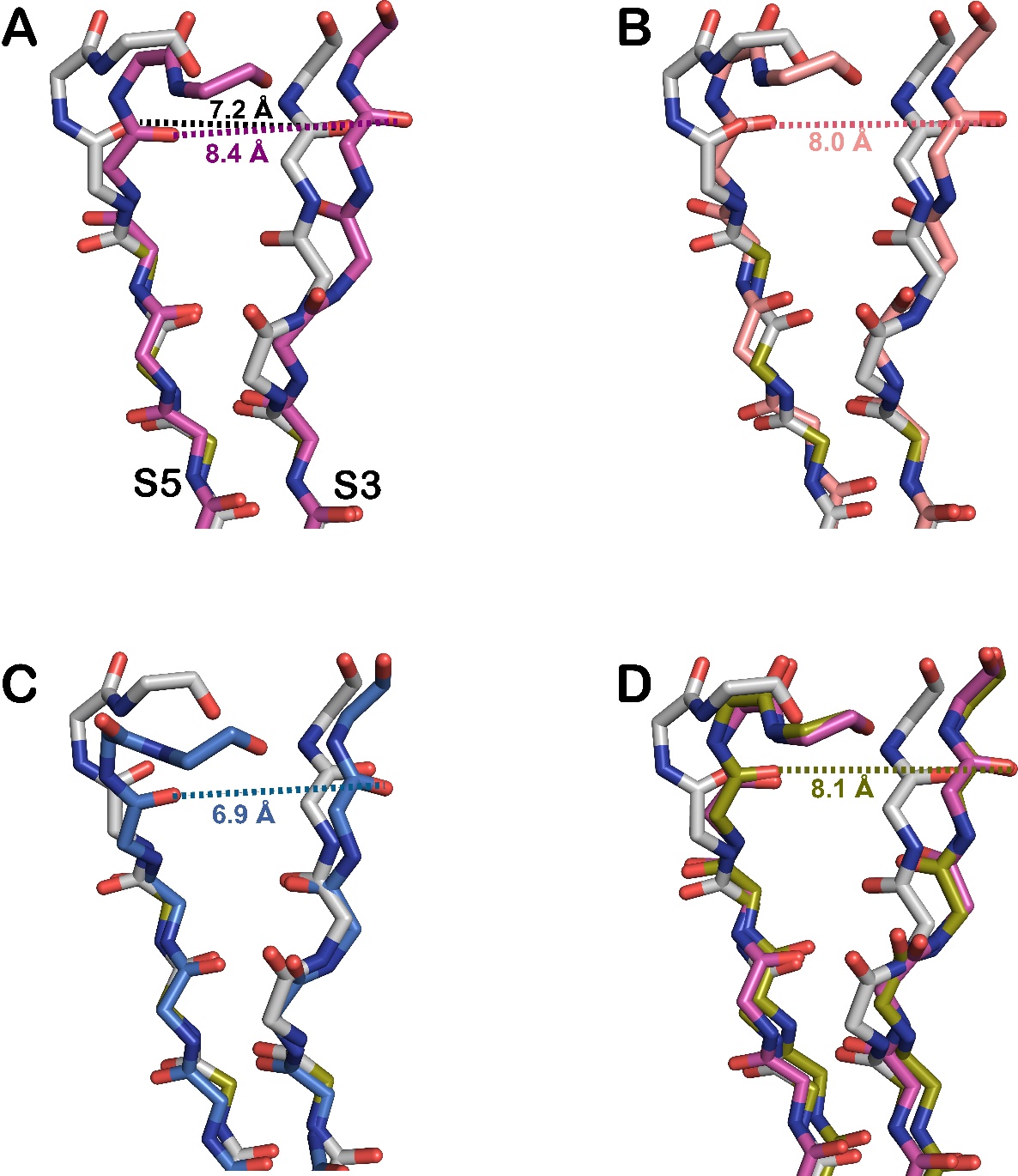

**Figure S1.** A slight strand separation between strands S3 and S5 of sheet A is seen between the WT and the mutants. Cartoon showing a superposition of the two strands between WT (grey) and A) E371K (magenta), B) E371R (pink), C) E371Q (blue) and D) E371K (magenta) and K318E-E371K (green).

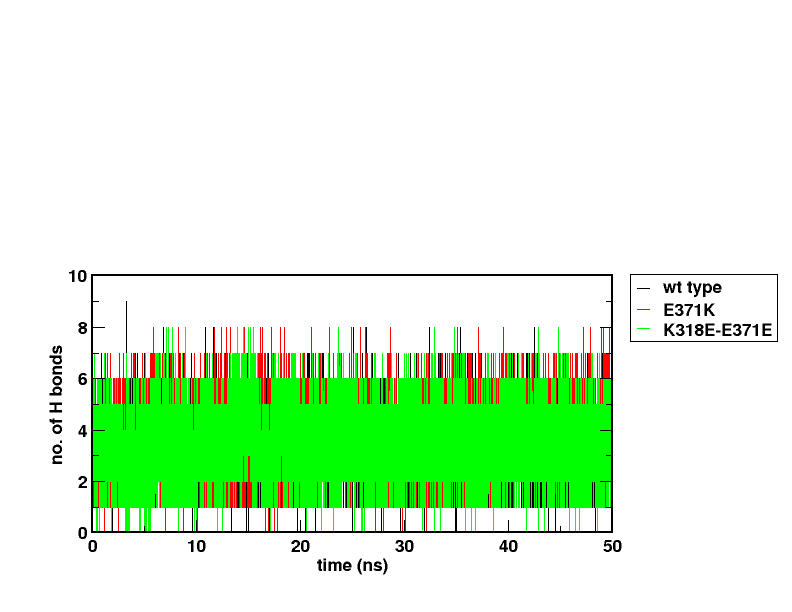

**Figure S2.** Inter-strand H bonds between S3A and S5A. Number of H-bonds between s3A and s5A main-chain atoms as a function of simulation time for WT, E371K and E371K-K318E variants (black, red, and green respectively).

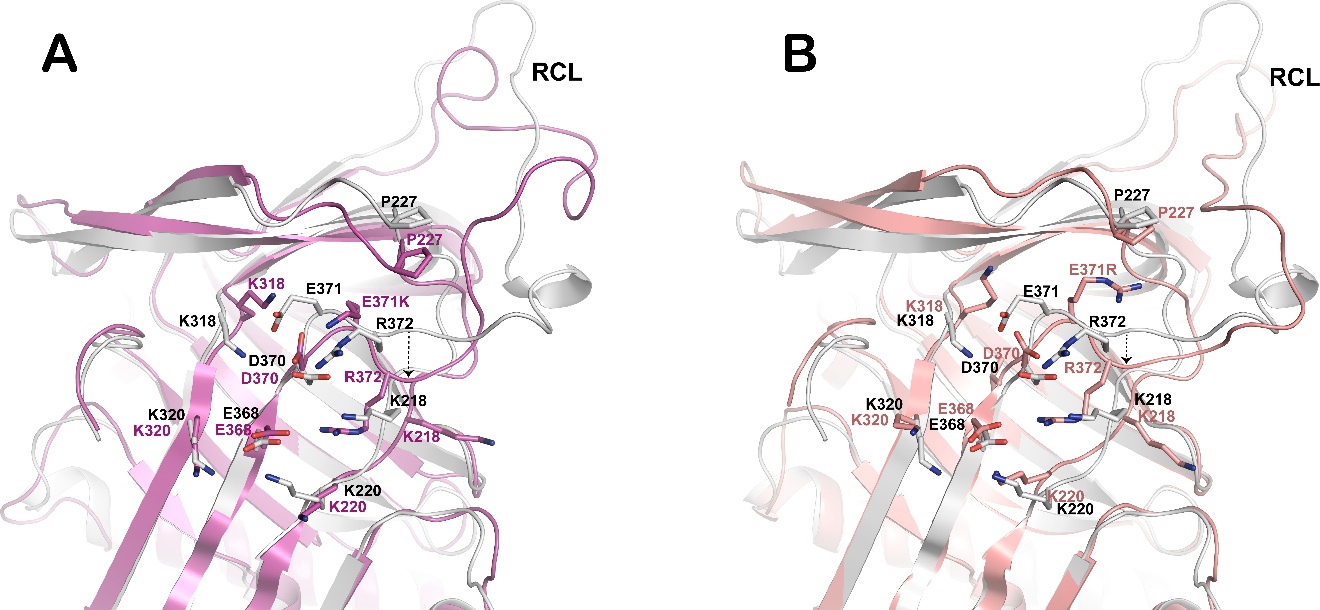

**Figure S3**. Superposition of the WT (grey) over (A) E371K (magenta) and (B) E371R (peach) shows insertion of the hinge region into the strand in the mutants. Note that in E371R, the E371R makes a stacking interaction with P227

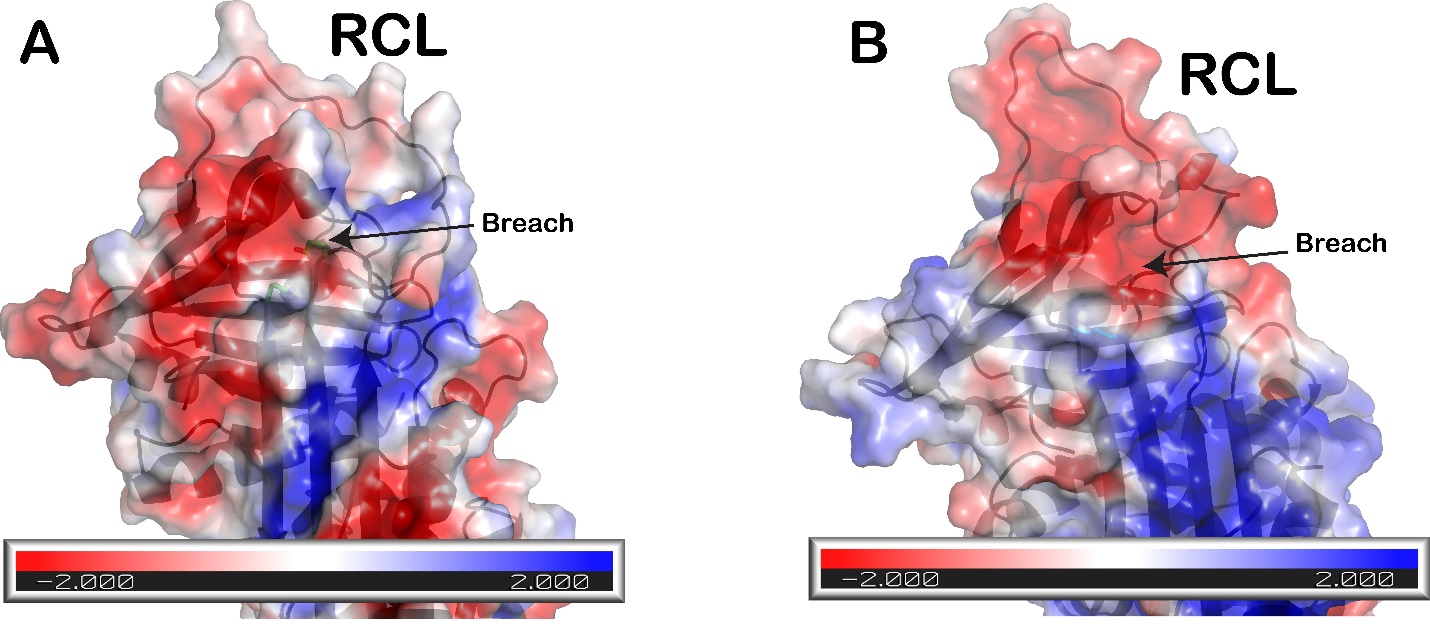

**Figure S4.** Surface electrostatic potentials of α1-AT (A) and ZPI (B) showing differences in the charge distribution near the breach region between the two proteins.
